# Compartmentalized glycolysis powers ATP production in primary cilia and engages mitochondria via the phosphoenolpyruvate cycle

**DOI:** 10.64898/2026.05.06.723011

**Authors:** Shih Ming Huang, Hannah R. Foster, Eun Young Lee, Jeong Hun Jo, Xinhang Dong, Byoung-Kyu Cho, Young Ah Goo, Jing W. Hughes, Matthew J. Merrins

## Abstract

Primary cilia are antenna-like sensory and signaling organelles present on most mammalian cells, including glucose-sensing pancreatic β-cells. Here, we show that the local energetic demands of primary cilia require the ATP-producing enzyme pyruvate kinase, with loss of PKm1, but not PKm2, impairing ciliary glycolytic flux. While the entire glycolytic machinery localizes to cilia, our data indicate that mitochondria are a critical source of phosphoenolpyruvate (PEP), the high-energy glycolytic intermediate that drives the pyruvate kinase reaction. Abolishing PCK2, the mitochondrial enzyme that generates PEP, prevents cilia from sensing not only glucose but also the amino acids glutamine and leucine. Finally, by mislocalizing glycolysis, we demonstrate that primary cilia can utilize ATP generated within the cell body when glucose is limiting. These findings indicate that primary cilia, while possessing the capacity for local ATP generation, leverage a ciliary-mitochondrial signaling axis to meet their bioenergetic needs.

## Introduction

Primary cilia are highly conserved, antenna-like sensory organelles that coordinate developmental, metabolic, and endocrine signaling pathways^1–7^. Despite lacking mitochondria, cilia carry out energetically demanding processes, including sensory transduction, receptor trafficking and motility^1,4,5,8–10^. Emerging evidence suggests that cilia possess intrinsic and local ATP-generating capacity. Glucose transporters and glycolytic enzymes have been detected within cilia of certain cell types, including neuroepithelial glial cells, photoreceptor rod cells, and olfactory neurons^11–18^. However, it remains unknown whether ATP is supplied through cytosolic diffusion or produced locally within the cilium. This distinction is critical, as reliance on distant mitochondrial ATP would tether ciliary signaling capacity to global cellular energy status, whereas if they could generate ATP autonomously, cilia-specific functions could be sustained independently of the cell body, with consequences for development, metabolism, and ciliopathy pathogenesis.

Pancreatic β-cells provide a powerful model to interrogate metabolic signaling because they are nutrient sensing cells with glycolytic and mitochondrial metabolism directly coupled to insulin secretion^19,20^. Their glucose sensing ability is partly due to glucokinase, a low-affinity hexokinase lacking product inhibition that enables high control of glycolytic flux across the physiological range of glucose concentrations^21^. However, unlike other metabolic cell types (e.g. muscle and liver) that maintain relatively stable ATP/ADP ratios, β-cells can raise the ATP/ADP ratio as a signaling mechanism to close K_ATP_ channels and initiate insulin secretion, even when their energetic needs are met^22–26^. Recent studies have demonstrated compartmentalized glycolytic regulation of the β-cell K_ATP_ channel by plasma membrane-associated glycolytic metabolons containing pyruvate kinase (PK), the enzyme that converts phosphoenolpyruvate (PEP) and ADP into ATP and pyruvate in the last step of glycolysis^19,27–29^. Compartmentalized glycolytic ATP production has also been found on mitochondria, ER/SR, and synaptic membranes in variety of cellular contexts^30–35^. These studies indicate that metabolism is not merely a bulk ATP-generating process but a spatially organized signaling network, raising the possibility that similar metabolic regulation operates within other specialized compartments, including the primary cilium.

As a specialized signaling organelle that lacks mitochondria, the primary cilium challenges us to understand energy regulation at subcellular resolution. Here, we combined three-dimensional live-cell imaging of cilia-targeted biosensors with spatial proteomics and genetic mouse models to define the metabolic architecture of β-cell cilia. By simultaneously monitoring the dynamics of glycolytic metabolites and ATP production in the cilia and cell body, we determined that cilia can locally generate ATP but are strongly dependent on extraciliary mitochondrial PEP production following a rise in glucose or amino acids. Our findings highlight metabolic compartmentalization as a key feature of cellular organization and define a mitochondria–cilium metabolic signaling axis that governs ciliary bioenergetics.

## Results

### Primary cilia are an autonomous metabolic compartment with ATP dynamics that are distinct from the plasma membrane

Primary cilia are externalized organelles that carry out compartmentalized trafficking and signaling processes requiring ATP. To investigate the dynamics of ciliary metabolism, we generated a β-cell cilia-targeted version of an ATP/ADP biosensor, Perceval-HR, by appending the ciliary targeting sequence from the 5-hydroxytryptamine (serotonin) receptor isoform 6 (5HT_6_)^36,37^. The sensor was expressed in mouse islet β-cells using adenovirus containing the insulin promoter. Three-dimensional live-cell imaging confirmed that 5HT_6_-Perceval-HR was enriched in the cilia and plasma membrane microvilli (Fig. 1A). To determine how glucose-stimulated ATP/ADP dynamics differ between the two compartments, we used machine learning– based tracking to resolve signals spatially and temporally (Fig. 1A, Supplemental Movie 1). Within a single islet, we observed synchronized plasma membrane ATP/ADP oscillations across β-cells (Fig. 1B). This coordinated behavior is well understood, and can be explained by gap junctions that facilitate intracellular Ca^2+^ wave propagation across the islet^38^. As the wave propagates, Ca^2+^ entry activates plasma membrane pumps that consume ATP^19^, which we detected as periodic lowering of plasma membrane ATP/ADP. In contrast to the plasma membrane, cilia displayed more heterogeneous ATP/ADP dynamics, with spontaneous ciliary ATP/ADP flashes of varying duration observed throughout the time course (Fig. 1B), an effect that was observable in all 25 islets tested (Fig. 1C). These ciliary ATP flashes were not noise artifacts, as their amplitude was nearly twice that of the plasma membrane (Fig. 1D), despite having a similar average frequency and duty cycle (Fig. 1E-F). Importantly, however, we observed no strong phase-locked relationship between plasma membrane and ciliary ATP dynamics (Fig. 1B,C), consistent with cilia functioning as autonomous metabolic compartments that consume ATP at rates distinct from the plasma membrane. Ciliary ATP dynamics were not asynchronous in all cases, as a subset of cilia intermittently exhibited a dominant pulse frequency that matched the cell body (Fig. 1G and Supplemental Fig. 1A-C), suggesting that under certain conditions, cilia may entrain to global cellular metabolic rhythms.

**Fig. 1:**
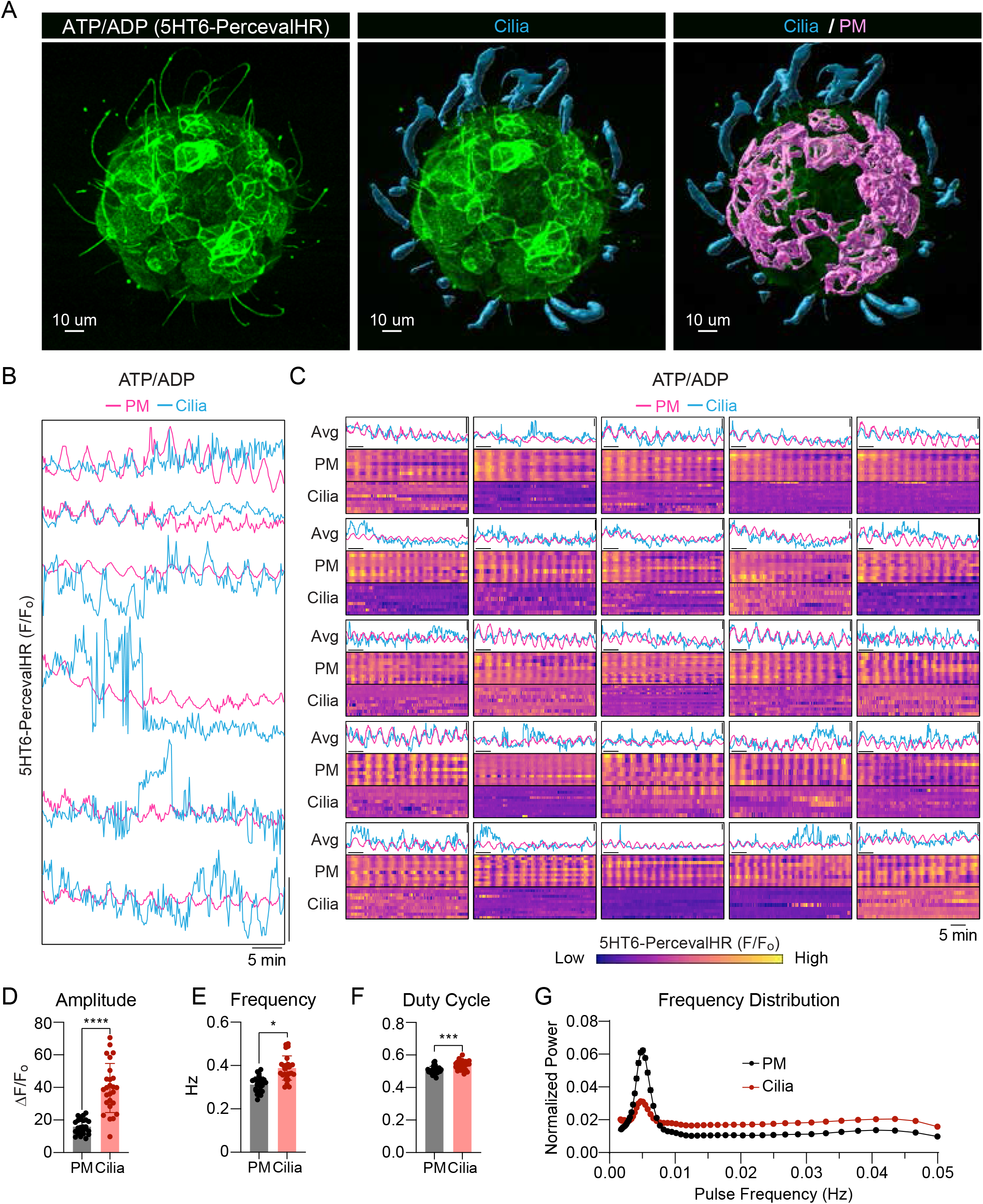
Primary cilia are autonomous metabolic compartments with distinct ATP/ADP dynamics from the plasma membrane in mouse islet β-cells. **A**, Three-dimensional image of a mouse islet β-cells expressing the cilia-targeted ATP/ADP biosensor 5HT_6_-Perceval-HR (green). Machine learning–based image segmentation was used to identify primary cilia (cyan) and the plasma membrane (magenta). Scale bar, 10 μm. **B**, Representative ATP/ADP traces at the plasma membrane (PM, magenta) and primary cilia (cyan) in mouse islet β-cells stimulated with 10 mM glucose. Overlaid traces represent paired measurements from the same β-cell, and all cells shown were recorded from a single islet. **C**, Heatmap representation of PM and ciliary ATP/ADP dynamics across 25 islets, with individual β-cells showing temporal activity in plasma membrane (PM) and ciliary compartments measured with 5HT_6_-PercevalHR under 10 mM glucose. Each row represents an individual β-cell, with paired PM and ciliary signals from the same cell. Traces above each heatmap show the corresponding average signal for each compartment of that islet. Color scale indicates normalized fluorescence (F/F_0_). **D–G**, Quantification of ATP/ADP oscillatory parameters comparing plasma membrane and cilia, including amplitude (**D**), frequency (**E**), duty cycle (**F**), and frequency distribution (**G**) (*n* =260 β-cells from 26 islets across 3 mice). Data are presented as individual islets with mean ± SEM. Statistical significance between PM and cilia was determined using *t*-test. *P* < 0.05, \*\**P* < 0.001, \*\*\**P* < 0.0001 **(D–G)**.

### Spatial proteomics and immunofluorescence imaging identifies glycolytic enzymes in human and mouse β-cell cilia

Primary cilia are approximately 5 μm in length but only 200 nm in diameter^39^, making them too small to accommodate mitochondria (Fig. 2A). Therefore, we looked for evidence of glycolytic machinery within primary cilia that could locally generate ATP in the absence of oxidative phosphorylation, using immunofluorescence to detect the subcellular localization of glucose transporters and glycolytic enzymes. First, we identified the presence of glucose transporters in cilia, including GLUT1 in human islets and GLUT2 in mouse islets, suggesting that cilia have the ability to locally sense glucose (Fig. 2B,C and Supplemental Fig. 2A,B). Within upper glycolysis, we further identified the presence of the rate-limiting enzyme glucokinase (GCK) in both human and mouse islet cilia (Fig. 2B,C). GCK, by catalyzing the conversion of glucose to glucose-6-phosphate, commits glucose carbon to glycolysis, while lower glycolysis is rate-limited by phosphofructokinase (PFK), which was also present (Fig. 2B,C). Although glyceraldehyde-3-phosphate dehydrogenase (GAPDH), the enzyme that converts NAD^+^ to NADH, was abundantly expressed in the cytosol of β-cells, it was also detected within primary cilia (Fig. 2B,C). β-cells express three isoforms of pyruvate kinase—PKm1, PKm2, and PKL^40^—with PKm1 and PKm2 accounting for the majority of enzymatic activity^28^. Immunostaining of mouse and human islets demonstrated ciliary localization of both PKm1 and PKm2. β-cell specific deletion of PKm1 (PKm1-βKO) and PKm2 (PKm2-βKO) confirmed the fidelity of the antibody staining (Supplemental Fig. 2C,D). Interestingly, β-cell cilia contain the LDHA isoform of lactate dehydrogenase (Supplemental Fig. 2E), which is lower in β-cells than many other metabolic cells such as myocytes, hepatocytes, and α-cells^29,41^, and would allow ciliary glycolysis to run at a high rate by locally regenerating NAD^+^ for GAPDH.

**Fig. 2:**
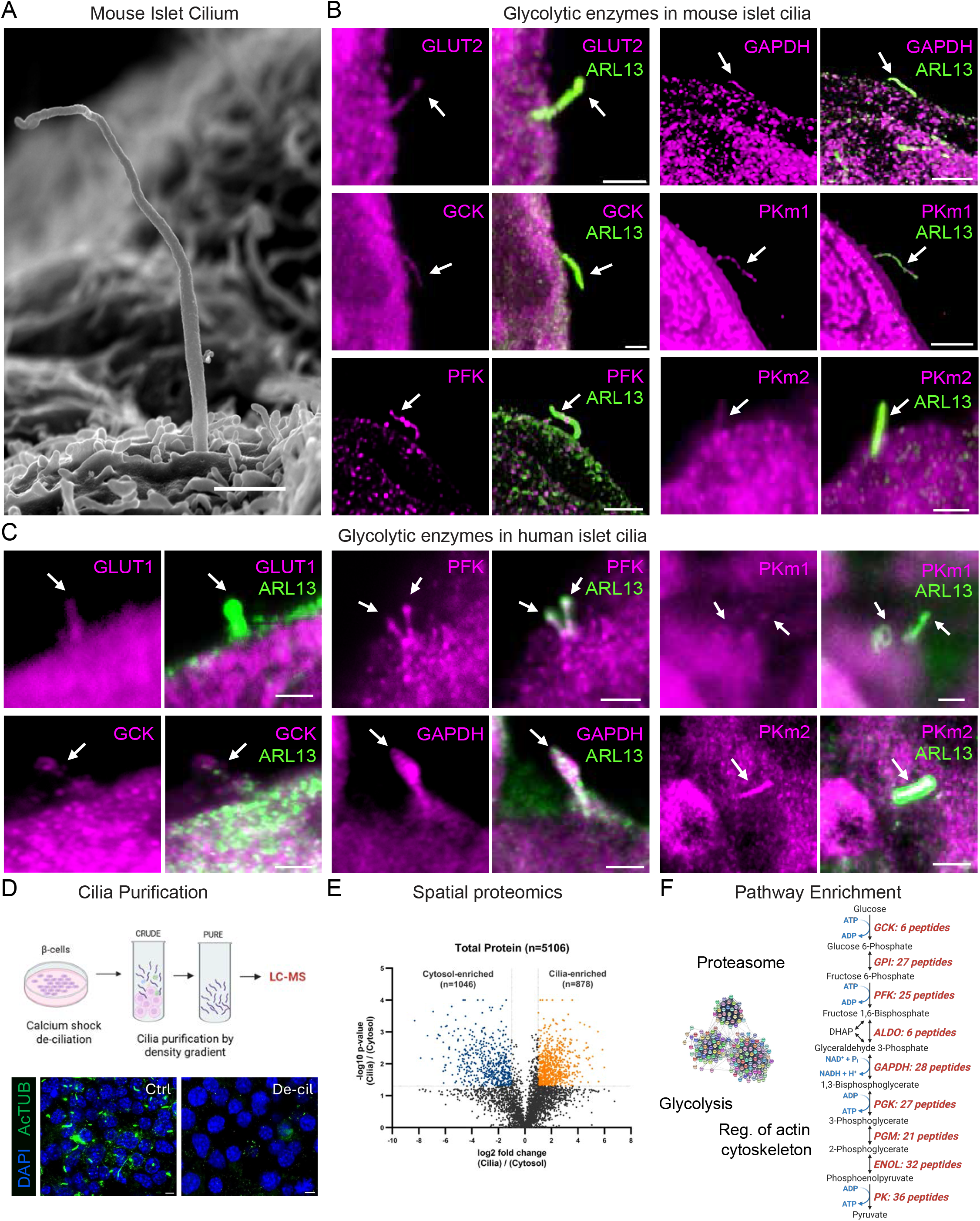
Immunofluorescence and proteomic detection of glycolytic enzymes in primary cilia. **A**, Scanning electron micrograph of a primary cilium protruding from the surface of a mouse pancreatic β-cell. Scale bar, 1 μm. **B**, Immunofluorescence images showing endogenous localization of glycolytic enzymes in mouse islet cilia. Glycolytic enzymes (magenta) are shown together with ciliary marker ARL13 (green). Arrows indicate cilia. Scale bar, 2 μm. **C**, Immunofluorescence images demonstrating endogenous localization of glycolytic proteins (magenta) in human islet cilia. Ciliary marker ARL13 (green). Arrows indicate cilia. Scale bar, 2 μm. **D**, Schematic of cilia purification workflow. Biochemical deciliation of mouse MIN6 β-cells was performed by calcium shock, followed by serial centrifugation to isolate cilia-enriched and cytoplasmic fractions for LC-MS/MS analysis. **E**, Spatial proteomics analysis comparing protein abundance between cytosolic and cilia-enriched fractions. Volcano plot shows enrichment of cilia-associated proteins in the isolated ciliary fraction. **F**, KEGG pathway enrichment analysis of the ciliary proteome highlighting glycolysis and actin cytoskeleton regulation among the top enriched categories, with the complete glycolytic enzyme chain identified.

To confirm the presence of glycolytic enzymes, we performed spatial proteomics on primary cilia isolated from MIN6 β-cells (Fig. 2D). Western blot confirmed efficient separation of primary cilia from the cell body in the isolated fractions (Supplemental Fig. 2F). Mass spectrometry identified a distinct and enriched ciliary proteome, including robust enrichment of canonical ciliary proteins, confirming successful compartment isolation and minimal contamination from non-ciliary cellular components (Fig. 2E). Comparative analysis against cytosol lysates demonstrated significant enrichment of proteins associated with the regulations of actin cytoskeleton and other cilia-related structural and signaling pathways. Importantly, all enzymes of the glycolytic pathway were identified within the ciliary fraction, by pathway enrichment analysis, including the key rate-limiting enzymes previously detected by immunofluorescence (Fig. 2F). Collectively, these data demonstrate that primary cilia of both mouse and human β-cells not only express the critical ATP-producing enzymes PKm1 and PKm2, but all the glycolytic machinery.

### The PKm1 isoform of pyruvate kinase is required to increase ATP/ADP in β-cell cilia, while PKm2 is dispensable

After establishing the localization of glycolytic enzymes in primary cilia of β-cell, we next sought to determine how ciliary bioenergetics are regulated. To monitor ciliary fluxes of glycolytic metabolites and signaling intermediates, we again used 5HT_6_-Perceval-HR to monitor the dynamics of ATP/ADP in the cilia and plasma membrane compartments. We also generated β-cell cilia–targeted biosensors for pyruvate (5HT_6_-PyronicSF) and fructose-1,6-bisphosphate (FBP, 5HT_6_-HYlight). Spinning-disk confocal imaging of isolated mouse islets expressing each biosensor confirmed the subcellular localization, demonstrating enrichment at both the cilium and plasma membrane (Fig. 3A,C,E). Collectively, these three biosensors are well-positioned to understand the activity of pyruvate kinase, since FBP is an allosteric activator of the PKm2 and PKL isoforms, while ATP and pyruvate are products of the reaction (Supplemental Fig. 3A).

**Fig. 3:**
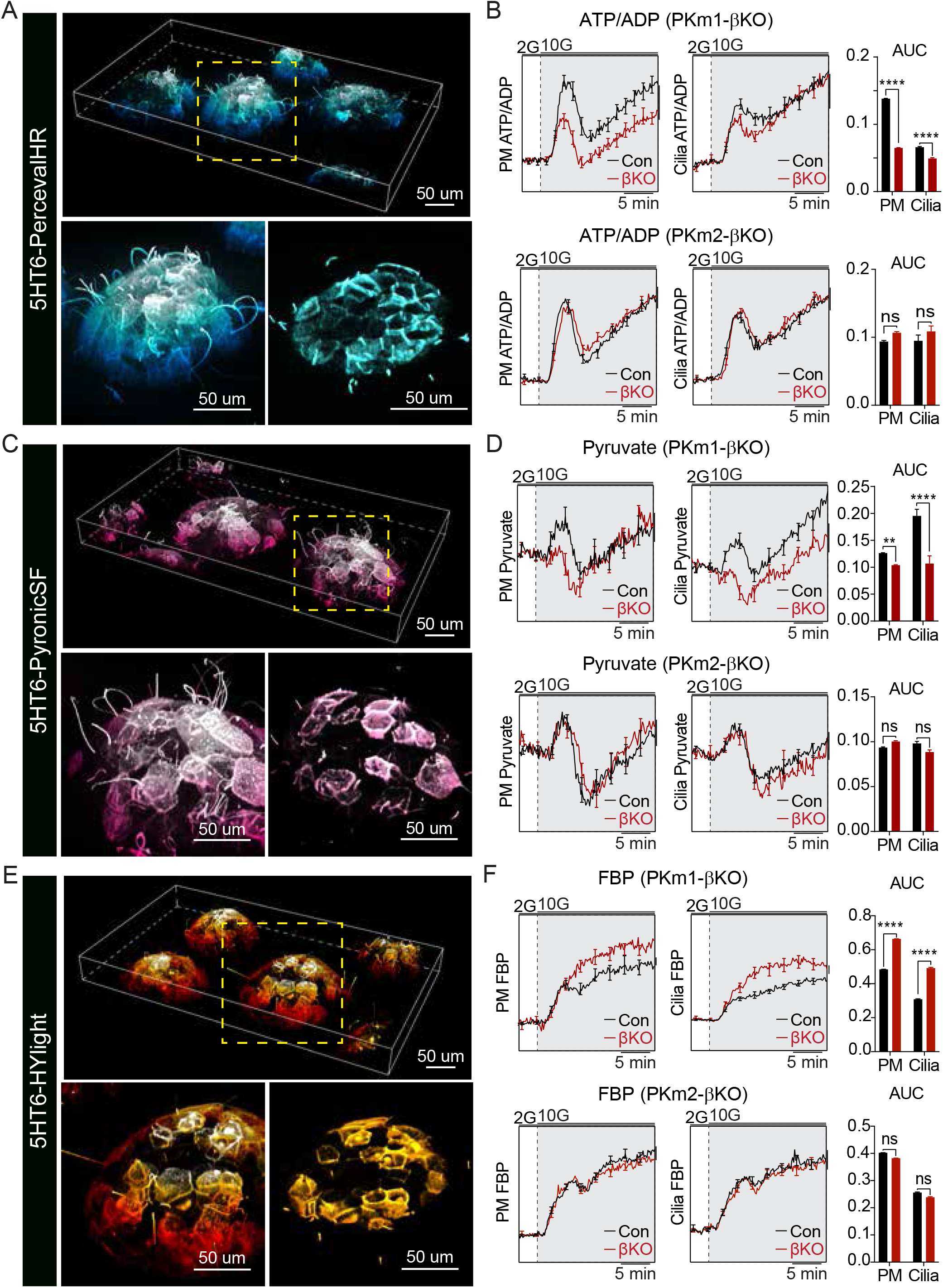
PKm1, but not PKm2, is required for glucose-stimulated glycolytic flux and ATP production in β-cell cilia. **A**,**C**,**E**, Representative three-dimensional images of mouse islets expressing β-cell cilia-targeted biosensors for ATP/ADP (5HT_6_-PercevalHR; **A**), pyruvate (5HT_6_-PyronicSF; **C**), and fructose-1,6-bisphosphate (FBP; 5HT_6_-HYLight; **E**). Insets show higher magnification of the boxed regions highlighting ciliary and plasma membrane localization of the biosensors. Scale bars, 50 μm. **B**,**D**,**F**, Dynamics of ATP/ADP (**B**), pyruvate (**D**), and FBP (**F**) measured in control, PKm1-βKO, and PKm2-βKO islets following glucose stimulation from 2 mM glucose (2G) to 10 mM glucose (10G) in the presence of 0.5 mM glutamine. Traces show average responses from plasma membrane (PM) and cilia, with quantification of the area under the curve (AUC) from PKm1-βKO (B, *n* = 211 cilia from 20 islets across 3 mice; D, *n* = 273 cilia from 25 islets across 4 mice; F, *n* = 164 cilia from 17 islets across 3 mice) and littermate controls (B, *n* = 253 cilia from 22 islets across 3 mice; D, *n* = 209 cilia from 18 islets across 4 mice; F, *n* = 207 cilia from 20 islets across 3 mice); PKm2-βKO (B, n=246 cilia from 22 islets across 3 mice; D, *n* = 155 cilia from 16 islets across 3 mice; F, n=239 cilia from 21 islets across 4 mice) and littermate controls (B, *n* = 280 cilia from 23 islets across 3 mice; D, *n* = 196 cilia from 18 islets across 3 mice; F, *n* = 218 cilia from 18 islets across 4 mice) Data are presented as mean ± SEM. Statistical significance was determined by t-test. **P < 0.01, ****P < 0.0001; ns, not significant.

In control islets, glucose stimulation elicited an initial rise in ciliary ATP/ADP followed by a sustained second phase (Fig. 3B). The plasma membrane exhibited similar behavior, with a more pronounced drop in ATP/ADP between the first and second phase. This drop coincides with β-cell Ca^2+^ influx (Supplemental Fig. 3B) and is likely due to consumption of ATP by Ca^2+^-ATPases^19^. Consistent with this interpretation, reducing Ca^2+^-driven workload with the application of cyclopiazonic acid (inhibits Ca^2+^-ATPases) and diazoxide (opens K_ATP_ channels to inhibit Ca^2+^ influx) increased the ATP/ADP ratio in the cilia and plasma membrane compartments by 102% and 403%, respectively (Supplemental Fig. 3B).

To determine the isoforms of pyruvate kinase that contribute to ciliary ATP production, we measured ATP/ADP dynamics in control and β-cell–specific PKm1 or PKm2 knockout mouse islets (PKm1-βKO and PKm2-βKO). In PKm1-βKO islets, the glucose-induced rise in ATP/ADP was markedly blunted in both the plasma membrane and ciliary compartments (Fig. 3B). While the plasma membrane ATP/ADP response remained persistently reduced in PKm1-βKO cells, ciliary ATP/ADP recovered to control levels during the second phase (Fig. 3B). These findings could reflect more efficient recruitment of PKm2 within the ciliary compartment (e.g., by FBP), or the lower workload in cilia (Supplemental Fig. 3B). However, deletion of β-cell PKm2 did not significantly alter ATP/ADP dynamics in either compartment, indicating that PKm1 is dominant for glucose-dependent ATP production within cilia (Fig. 3B).

The concentration of pyruvate in cilia is determined by the activity of pyruvate kinase, consumption by lactate dehydrogenase, as well as potential efflux to the cell body. Following glucose stimulation, PKm1-βKO islets displayed a markedly attenuated pyruvate rise at both the plasma membrane and cilia (Fig. 3D). Moreover, ciliary pyruvate levels failed to return to control levels over time (Fig. 3D). Consistent with reduced ATP and pyruvate generation, the upstream metabolite FBP accumulated in PKm1-βKO upon glucose stimulation (Fig. 3F), reflecting a metabolic bottleneck at the pyruvate kinase step. Although PKm2 is present in cilia (Fig. 1B), PKm2 deletion did not significantly alter either pyruvate or FBP responses (Fig. 3D,F). Together, these findings demonstrate that PKm1 is required for efficient glycolytic flux and glucose-stimulated ATP production in both the plasma membrane and primary cilium, establishing PKm1 as the essential isoform regulating β-cell cilia bioenergetics.

### Glucose fuels a ciliary-mitochondrial PEP cycle

In the PEP cycle, pyruvate generated by glycolysis is first metabolized by mitochondria pyruvate carboxylase to oxaloacetate, and then converted by mitochondrial phosphoenolpyruvate carboxykinase (PCK2) to PEP, which is exported to the cytosol. Our prior studies have shown that mitochondrially-derived PEP contributes to pyruvate kinase activity at the plasma membrane by supplementing the PEP provided directly by glycolysis^28^. However, these studies, which utilized β-cell PCK2 deletion (PCK2-βKO), were conducted in the absence of glutamine, which we later found to be essential for the PEP cycle in pancreatic α-cells^42^. To test whether β-cells have a similar requirement for glutamine, it was necessary to reexamine cytosolic ATP/ADP dynamics in PCK2-βKO islets in the presence of 0.5 mM glutamine (which was used throughout this paper). Islets from knockout and littermate control mice were simultaneously measured using a far-red dye (DiD) barcoding system (Fig. 4B) that does not impact islet function^28,42^. The glucose-dependent rise of cytosolic ATP/ADP was reduced by 35% in PCK2-βKO islets relative to controls (Fig. 4B), and insulin secretion was reduced by 30% (Supplemental Fig. 4A). This defect was not due to an impairment of the proton motive force, which powers mitochondrial ATP production by oxidative phosphorylation, as mitochondrial membrane potential was unaffected by PCK2 deletion in the presence of elevated glucose (Fig. 4C). These data indicate that the phenotype of β-cell PCK2 knockout is more severe than previously reported^28^, as mitochondrial PEP production is required for cytosolic ATP/ADP production and insulin secretion in the presence of both glucose and glutamine.

**Fig. 4:**
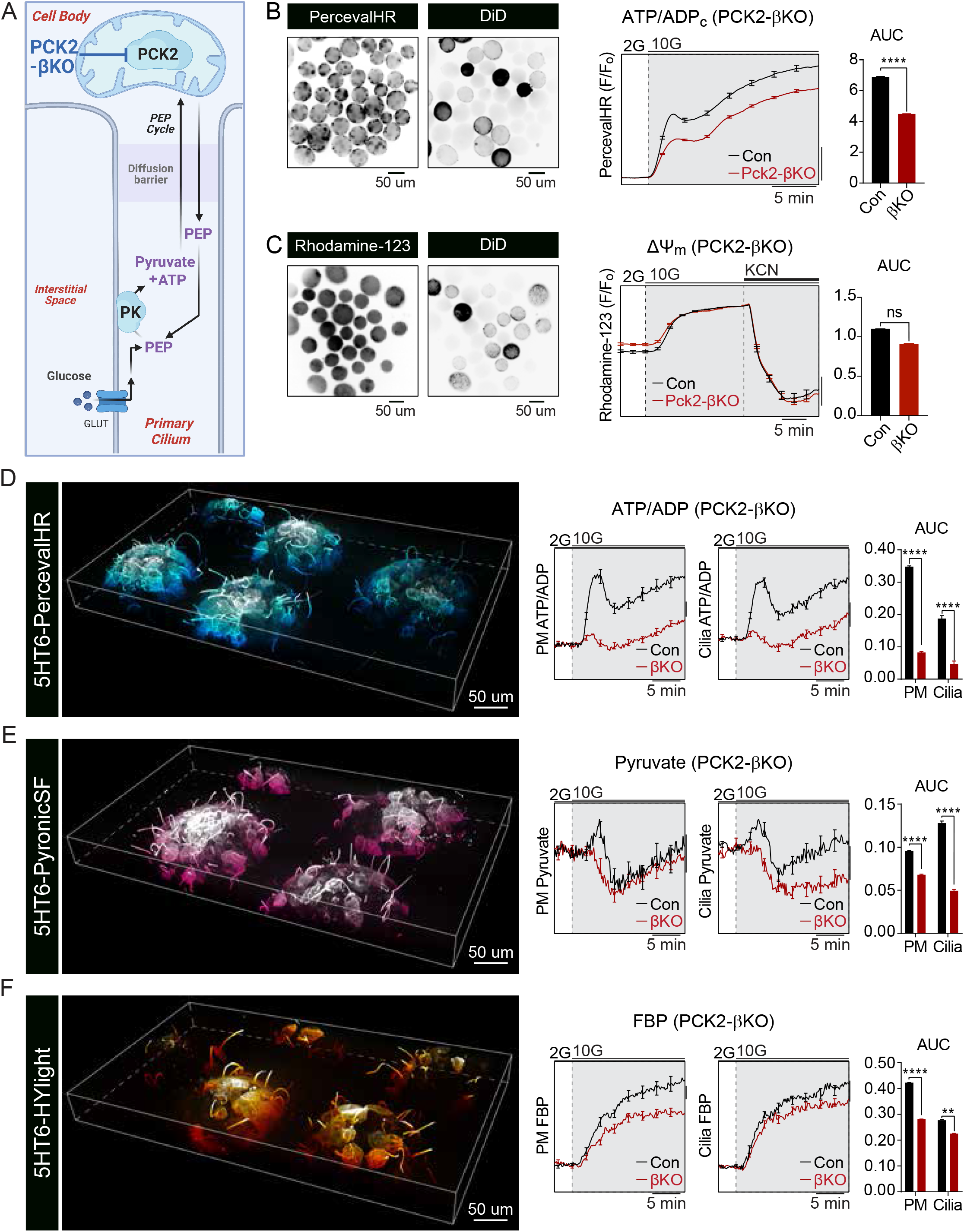
The mitochondrial PEP cycle, mediated by PCK2, regulates glucose-stimulated glycolytic flux and ATP production in β-cell cilia. **A**, Proposed model linking the mitochondrial phosphoenolpyruvate (PEP) cycle with ciliary bioenergetics. Mitochondrial PCK2 generates PEP that supplies pyruvate kinase (PK) to produce pyruvate and ATP within the ciliary compartment. **B**,**C**, Cytosolic ATP/ADP (ATP/ADP_c_) responses (**B**) and mitochondrial membrane potential (ΔΨ_m_) measured in control and PCK2-βKO islets during glucose stimulation from 2 mM to 10 mM glucose in the presence of 0.5 mM glutamine. For mitochondrial measurements (**C**), the complex IV inhibitor cyanide (KCN, 3 mM) was subsequently applied to induce mitochondrial depolarization. Islets from different genotypes were distinguished by DiD barcoding. ATP/ADP_c_ responses were measured using Perceval-HR and quantified as area under the curve (AUC) from PCK2-βKO (*n* = 103 islets from 3 mice) and littermate controls (*n* = 118 islets from 3 mice). Mitochondrial membrane potential was measured using Rhodamine-123, normalized to fluorescence after KCN-induced depolarization, and quantified as AUC for 10G treatment from PCK2-βKO (*n* = 76 islets from 3 mice) and littermate controls (*n* = 82 islets from 3 mice). **D–F**, Three-dimensional images of mouse islets expressing cilia-targeted biosensors for ATP/ADP (5HT_6_-PercevalHR; **D**), pyruvate (5HT_6_-PyronicSF; **E**), and fructose-1,6-bisphosphate (FBP; 5HT_6_-HYLight; **F**). Corresponding traces show responses measured at the plasma membrane (PM) and cilia in control and PCK2-βKO islets following glucose stimulation from 2 mM to 10 mM glucose in the presence of 0.5 mM glutamine. Bar graphs show quantification of AUC from PCK2-βKO (D, *n* = 239 cilia from 21 islets across 3 mice; E, *n* = 267 cilia from 21 islets across 3 mice; F, *n* = 180 cilia from 16 islets across 3 mice) and littermate controls (D, *n* = 238 cilia from 21 islets across 3 mice; E, *n* = 254 cilia from 17 islets across 3 mice; F, *n* = 266 cilia from 23 islets across 3 mice). Data are presented as mean ± SEM. Statistical significance was determined by t-test. **P < 0.01, ****P < 0.0001; ns, not significant.

If ciliary PK communicates with mitochondria similarly to the plasma membrane, we reasoned that β-cell deletion of PCK2 would compromise local ATP production in cilia. We therefore compared glucose-stimulated ATP/ADP, pyruvate, and FBP dynamics at the plasma membrane and primary cilia in PCK2-βKO islets. Following glucose elevation, PCK2-βKO islets exhibited significantly attenuated responses across all three metabolites in both compartments relative to controls, with ciliary ATP/ADP responses reduced by >75% (Fig. 4D-F). Glucose-stimulated pyruvate dynamics were also markedly blunted at both the plasma membrane and cilia, consistent with impaired pyruvate kinase activity downstream of mitochondrial PEP production (Fig. 4E). Notably, while the plasma membrane pyruvate level recovered over time, cilia pyruvate did not. Similarly, FBP responses were reduced in both compartments, indicating decreased flux through upper glycolysis (Fig. 4F). These findings indicate that ciliary ATP production is dependent on mitochondrial PEP generation, and does so to a greater extent than the cell body.

### Cilia sense amino acids via the mitochondrial production of PEP

To determine whether mitochondria supply PEP to fuel ciliary ATP production, we stimulated β-cells with the mitochondrial fuels leucine and glutamine. Leucine, the only amino acid sufficient to stimulate insulin secretion, activates glutamate dehydrogenase to stimulate the glutamine-dependent production of mitochondrial PEP, which in turn fuels ATP generation via pyruvate kinase^43–45^. To minimize glycolytic PEP production, these experiments were conducted at low glucose (2 mM, well below the K_m_ of glucokinase), thereby isolating mitochondria as the primary source of PEP. Under these conditions, PCK2-βKO islets displayed profound defects in amino acid–stimulated metabolism. ATP/ADP responses were significantly reduced at both the plasma membrane and cilia, and were accompanied by marked attenuation of pyruvate and FBP in both compartments (Fig. 5B-D). As expected, when glycolytic PEP is limited, loss of PCK2 disrupted the mitochondrial PEP supply to pyruvate kinase, impairing ATP production. Consistent with these findings, leucine failed to stimulate insulin secretion in both PCK2-βKO and PKm1-βKO islets (Supplementary Fig. 5A,B), linking the PEP-dependent ATP deficit to impaired secretory function.

**Fig. 5:**
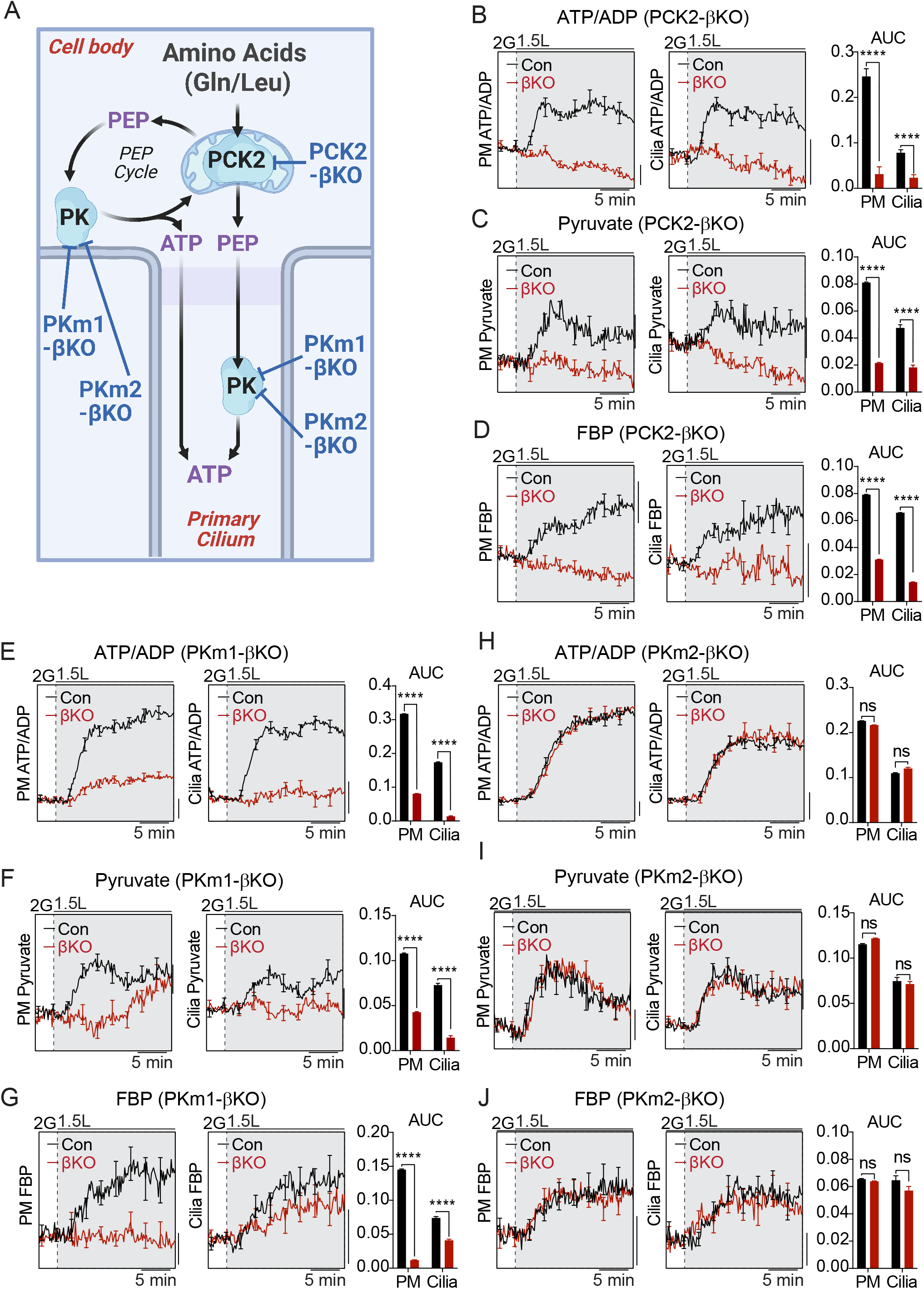
Retrograde mitochondria–cilium metabolic signaling regulates ATP production in β-cell cilia. **A**, Schematic illustrating a hypothesized model for retrograde metabolic communication between mitochondria and the primary cilium. Leucine and glutamine fuels mitochondrial PEP production via PCK2. In the first route, mitochondrial PEP is supplied directly to pyruvate kinase located within the cilium to support local ATP generation. In the second route, mitochondrial PEP is first utilized by plasma membrane PK to generate ATP, which subsequently diffuses into the cilium to support its energetic demand. **B–J**, Dynamics of ATP/ADP (**B**,**E**,**H**), pyruvate (**C**,**F**,**I**), and FBP (**D**,**G**,**J**) measured at the plasma membrane (PM) and cilia in control, PCK2-βKO (**B–D**), PKm1-βKO (**E–G**), and PKm2-βKO (**H–J**) islets following stimulation with leucine (1.5 mM) at low glucose (2 mM) in the presence of 0.5 mM glutamine. Average traces are shown with quantification of the area under the curve (AUC) from PCK2-βKO (B, *n* = 212 cilia from 14 islets across 3 mice; C, *n* = 233 cilia from 20 islets across 3 mice; D, *n* = 201 cilia from 20 islets across 3 mice) and littermate controls (B, *n* = 237 cilia from 14 islets across 3 mice; C, *n* = 241 cilia from 20 islets across 3 mice; D, *n* = 241 cilia from 18 islets across 3 mice); PKm1-βKO (E, *n* = 223 cilia from 21 islets across 3 mice; F, *n* = 290 cilia from 29 islets across 4 mice; G, *n* = 202 cilia from 19 islets across 3 mice) and littermate controls (E, *n* = 233 cilia from 21 islets across 3 mice; F, *n* = 221 cilia from 23 islets across 4 mice; G, *n* = 165 cilia from 12 islets across 3 mice); PKm2-βKO (H, *n* = 255 cilia from 17 islets across 3 mice; I, *n* = 196 cilia from 17 islets across 3 mice; J, *n* = 232 cilia from 14 islets across 3 mice) and littermate controls (H, *n* = 203 cilia from 18 islets across 3 mice; I, *n* = 189 cilia from 14 islets across 3 mice; J, *n* = 201 cilia from 11 islets across 3 mice) Data are presented as mean ± SEM Statistical significance was determined by t-test.

Similarly, in PKm1-βKO islets, amino acid stimulation elicited markedly blunted ATP/ADP responses at the plasma membrane and within cilia as compared to controls (Fig. 5E). The rise of pyruvate and FBP were also attenuated (Fig. 5F,G), indicating that PKm1 is required for cilia to mount a metabolic response to glutamine/leucine. In contrast, PKm2-βKO islets exhibited no significant differences in metabolic responses to amino acids. ATP/ADP, pyruvate, and FBP dynamics at both compartments were comparable to controls (Fig. 5H-J), supporting previous findings that PKm2 is not essential for ciliary ATP production. These findings demonstrate that mitochondrial PEP production via PCK2, and its utilization by PKm1, are required for amino acid-stimulated ATP generation in cilia.

### Plasma membrane ATP production can rescue ciliary ATP/ADP responses under low-glucose conditions

A diffusion barrier at the base of the primary cilium regulates molecular exchange between the ciliary compartment and the cell body^46,47^. Determining whether ATP can cross this barrier— or must be generated locally—is essential to understand whether primary cilia function as energetically autonomous signaling domains. To address this, we took advantage of the bioenergetic defects in β-cells lacking PKm1 and then re-introduced PKm1 targeted to different subcellular compartments. First, we generated an expression construct in which PKm1 was targeted to the plasma membrane as well as the primary cilium (5HT_6_-mCherry-PKm1), as confirmed by co-localization with 5HT_6_-Percerval-HR (Fig. 6A). In contrast, AKAP79-mCherry-PKm1 exhibited strong and selective localization to the plasma membrane, with no detectable ciliary expression (Fig. 6C). Expression of 5HT_6_-mCherry-PKm1 significantly enhanced ciliary ATP/ADP responses under glucose stimulation compared to neighboring β-cells from the same islet lacking PKm1 (Fig. 6B). Strikingly, expression of plasma membrane-targeted AKAP79-mCherry-PKm1 restored ciliary ATP/ADP with kinetics that were indistinguishable from direct cilia targeting of PKm1 (Fig. 6D). As both constructs localized to the plasma membrane, and both rescued ciliary ATP/ADP responses, this finding demonstrates that PKm1 need not be intra-ciliary to support ciliary energetics; expression at the plasma membrane is sufficient. This experiment is consistent with free diffusion of ATP from the cell body into the cilium, and indicates that ATP generated at the plasma membrane is sufficient to support ciliary ATP/ADP responses when glucose is limiting. We note, however, that under persistently elevated glucose conditions, cilia function more independently and are not required to follow the ATP/ADP ratio in the cell body (Fig. 1). Collectively, these findings indicate that primary cilia can leverage ATP diffusion from the cell body under energy starved conditions, while functioning as an autonomous metabolic compartment when glucose is replete.

**Fig. 6:**
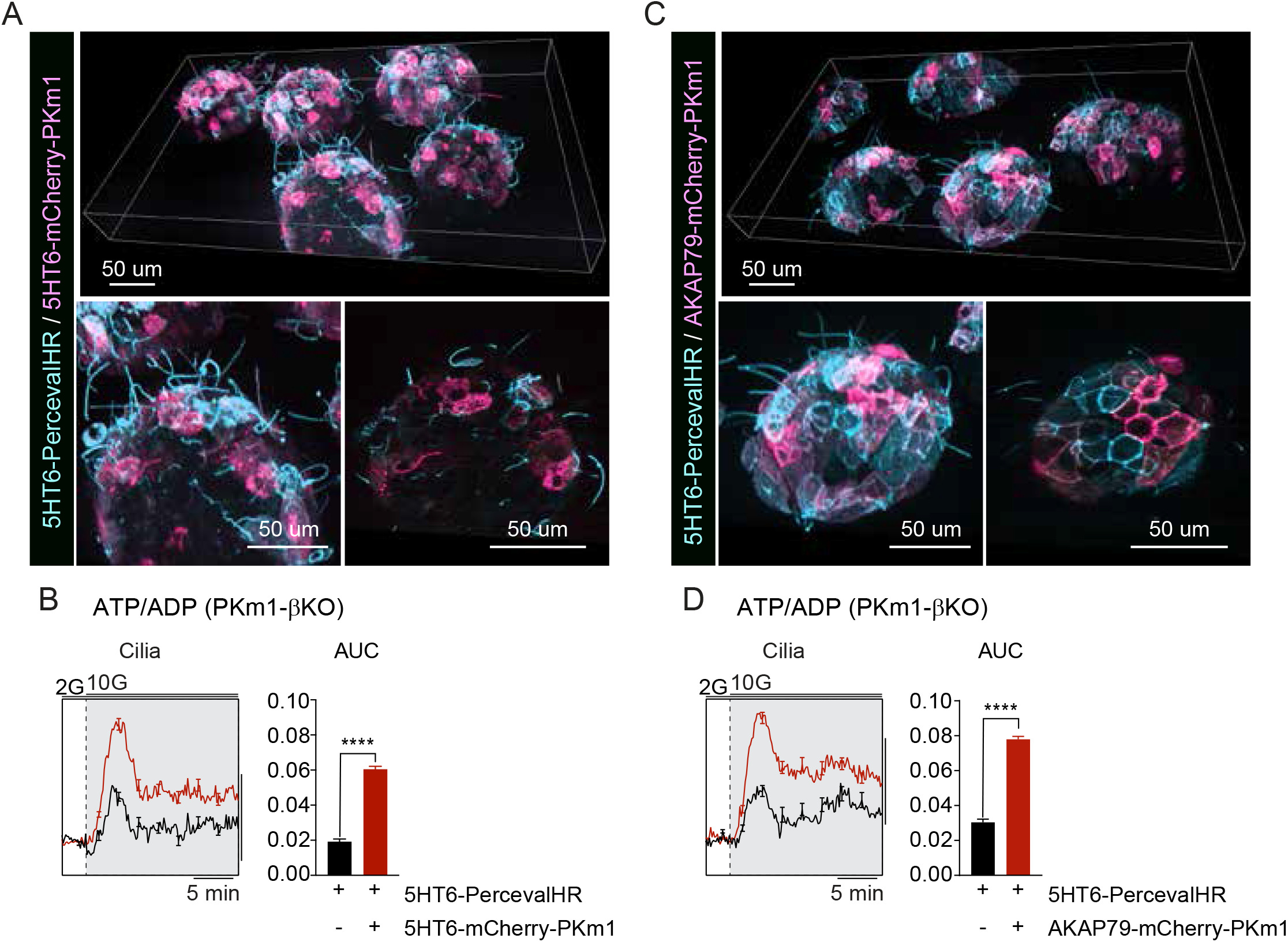
Plasma membrane–generated ATP supports ciliary energetics in β-cells. **A**,**C**, Representative 3D images of mouse islets expressing the ciliary-ATP/ADP biosensor 5HT_6_-Perceval-HR (cyan) together with PKm1 targeted either to the plasma membrane and primary cilium via the 5HT_6_ sequence (5HT_6_-mCherry-PKm1; **A**) or selectively to the plasma membrane via AKAP79 (AKAP79-mCherry-PKm1; **C**). Magenta indicates mCherry-PKm1. Insets show higher-magnification views highlighting subcellular localization. **B**,**D**, ATP/ADP dynamics measured using 5HT_6_-Perceval-HR in PKm1-βKO islets expressing 5HT_6_-mCherry-PKm1 (**B**) or AKAP79-mCherry-PKm1 (**D**) following glucose stimulation (2 mM to 10 mM glucose) in the presence of 0.5 mM glutamine. Average traces are shown with quantification of area under the curve comparing β-cells expressing both the PKm1 construct and the biosensor to neighboring β-cells expressing the biosensor alone. Measurements were collected from 47 islets for the 5HT_6_-PKm1 experiments (biosensor only: 235 cilia; PKm1-expressing: 282 cilia) and 46 islets for the AKAP79-PKm1 experiments (biosensor only: 227 cilia; PKm1-expressing: 233 cilia) from 5 PKm1-βKO mice.

## Discussion

In this study, we demonstrate that primary cilia in pancreatic β-cells are metabolically specialized signaling compartments with ATP dynamics distinct from the plasma membrane. Spatial proteomics revealed the presence of all the glycolytic enzymes within cilia, including both PKm1 and PKm2. Functional analysis identified PKm1 as the dominant isoform required for glucose- and amino acid–stimulated ATP production within cilia, whereas PKm2 was not essential. We further show that mitochondrial PEP production via PCK2 is essential for ciliary ATP generation, establishing retrograde metabolic communication from mitochondria to cilia. Together, these findings define a mitochondria–cilium metabolic signaling axis governing compartment-specific ATP dynamics.

Our data indicate that glycolysis within the primary cilium and at the plasma membrane is functionally distinct from the bulk cytosol. Previous work by Foster and colleagues showed that cytosolic ATP/ADP remains largely preserved in PKm1-βKO islets during glucose stimulation^28^, indicating that overall cellular ATP production in the cell body can buffer global energetic demand and maintain bulk ATP level. In contrast, we observe pronounced deficits in ATP dynamics specifically within ciliary and plasma membrane compartments in PKm1-βKO islets. The spatial divergence between the cell body and plasma membrane indicates that PKm1 activity is not required to maintain cytosolic ATP levels but is critical for supporting ATP production within the more energetically demanding plasma membrane domain. Thus, the metabolic workload differs substantially across compartments, with mitochondria-rich regions of the cell body providing robust ATP buffering capacity, whereas the plasma membrane experiences rapid ATP turnover due to ion channels and ATPases that support Ca^2+^ handling. The primary cilium represents an even more specialized compartment, lacking mitochondria entirely and therefore relying on localized glycolysis and substrate exchange with the cell body to meet its energetic demands. In this framework, the enrichment of glycolytic enzymes within cilia and the dependence on mitochondrial PEP cycling support a model in which spatially organized glycolytic metabolons are tailored to match compartment-specific workloads while complementing mitochondrial ATP production in the cell body^29–31^.

Mechanistically, our data support two parallel energetic routes linking mitochondria, plasma membrane, and cilium. In the first route, mitochondrial PEP produced by PCK2 is supplied directly to pyruvate kinase located at the plasma membrane and within the cilium, enabling local ATP generation at these compartments. In the second route, mitochondrial PEP is first utilized by plasma membrane pyruvate kinase to generate ATP, and the ATP produced subsequently diffuses into the cilium to support its energetic demand. Our glycolytic mis-localization experiments demonstrate that ATP produced at the plasma membrane can diffuse into the ciliary compartment. Whether PEP itself traverses the ciliary diffusion barrier remains unresolved, although it is likely an excellent signaling molecule because it has few interacting partners and contains a high free energy bond twice that of ATP. Pyruvate may also serve as a mobile intermediate due to its central position linking glycolysis, mitochondrial metabolism, and lactate production. The presence of LDHA within cilia further suggests that localized redox shuttling contributes to sustaining glycolytic flux, enabling rapid regeneration of NAD^+^ required for GAPDH activity. Together, these mechanisms suggest that ciliary energetics are sustained not by a single diffusible energy carrier but by coordinated exchange of metabolites and redox equivalents across subcellular compartments, reinforcing the concept that metabolic signaling microdomains organize energy flow within the β-cell^19,48^.

Our findings further suggest that primary cilia may function as sensors of mitochondrial function. Because ciliary ATP production depends strongly on mitochondrial PEP, perturbations in mitochondrial metabolism could impair ciliary signaling even when the cilium itself remains structurally intact. In this framework, defects in mitochondrial nutrient metabolism may propagate to the cilium through metabolic intermediates, altering ciliary ATP dynamics and thereby influencing energy-dependent signaling processes. Our amino acid stimulation experiments further support this concept by demonstrating that mitochondrial nutrient sensing directly regulates ciliary bioenergetics through the PEP–pyruvate kinase axis, an observation that likely explains how cilia sense glutamine deprivation on longer timescales^49^. These observations position the primary cilium downstream of mitochondrial nutrient sensing pathways, suggesting that ciliary signaling may integrate metabolic information from the cell body to coordinate cellular responses to nutrient availability. More broadly, these findings reveal how metabolic communication between organelles can shape the energetic landscape of signaling compartments and highlight metabolic compartmentalization as a fundamental principle linking mitochondrial function to ciliary signaling.

## Methods

### Mice and islet isolations

All experiments involving animals were conducted according to the National Institutes of Health’s *Guide for the Care and Use of Laboratory Animals* and approved by the Institutional Animal Care and Use Committees of the University of Wisconsin-Madison (Madison, WI), and Yale University (New Haven, CT). Wild-type C57BL/6J mice were obtained from The Jackson Laboratory. β-cell-specific PCK2, PKm1 and PKm2 knockout mice were generated as previously described^28^. Briefly, PCK2^*f/f*^ mice were generated de novo by the University of Wisconsin-Madison Genome Editing and Animal Model core and crossed with *Ucn3-Cre* mice^50^ (Tg(Ucn3-cre)KF43Gsat/Mmucd, MMRRC_032078-UCD). *Ins1-Cre* mice^51^ (B6(Cg)-*Ins1*^*tm1*.*1(cre)Thor*^*/J*, Jackson Laboratory #026801) was crossed with Pkm1^*f/f*^ mice^52,53^ or Pkm2^*f/f*^ mice^54^ (B6;129S-*Pkm*^*tm1*.*1Mgvh*^/J, Jackson Laboratory #024408) to generate PKm1-βKO and PKm2-βKO mice. Littermate *Ins1-Cre* mice were used as controls for PKm1-βKO and PKm2-βKO mice, while littermate *PCK2*^*f/f*^ mice were used as controls for PCK2-βKO mice. All mice were genotyped by Transnetyx. Male mice were housed 1-5 per cage at 21-23°C, maintained on a 12-hour light/dark cycle, with chow diet and water provided ad libitum. Male mice 12–18 weeks of age were sacrificed via CO_2_ asphyxiation followed by cervical dislocation, and islet isolations were carried out as previously described^55^.

### Cloning and adenoviral delivery of biosensors

Generation of adenovirus for β-cell specific expression of the ATP/ADP biosensor Perceval-HR was described previously^56^. Adenoviruses encoding 5HT_6_-Perceval-HR, the pyruvate sensor 5HT_6_-PyronicSF (Addgene #124812), and the FBP sensor 5HT6-HYLight (Addgene #193447)^57^ were commercially generated by VectorBuilder by cloning each construct into an adenovirus serotype 5 backbone under the rat insulin promoter with a rabbit β-globin intron and the 5HT_6_ ciliary targeting sequence (Addgene #47500)^37^ on the N-terminus of each biosensor. For rescue experiments, PKm1 was fused to mCherry and targeted either to the primary cilium using the 5HT_6_ targeting sequence (5HT_6_-mCherry-PKm1) or to the plasma membrane (AKAP79-mCherry-PKm1) using the membrane-targeting sequence from A-kinase anchoring protein 79 (AKAP79)^58,59^. Islets were infected immediately after isolation with high-titer adenovirus for 2 h at 37 °C, followed by overnight culture in fresh medium. Imaging experiments were performed 3 days post-infection.

### Live-cell imaging

For measurements of ciliary and plasma membrane ATP/ADP, pyruvate, and FBP, islets expressing corresponding biosensor were placed in a glass-bottomed imaging chamber (Warner Instruments) on Nikon CSU-W1 Spinning Disk Confocal Microscope with a Plan Apo 40x/1.25 NA silicone immersion objective (Nikon Instruments). The chamber was perfused with standard imaging solution (in mM: 135 NaCl, 4.8 KCl, 2.5 CaCl_2_, 1.2 MgCl_2,_ 20 HEPES, 0.5 glutamine) with glucose and amino acids concentrations indicated in the figure legends. Temperature was maintained at 33°C with an SF inline solution heater and QE-1 chamber heaters (Warner Instruments), and the flow rate was set to 0.25 mL/min. 5HT_6_-Perceval-HR, 5HT_6_-PyronicsSF, and 5HT_6_-HYlight were excited at 488 nm (1.5% power) and emission was collected with Hamamatsu Orca Quest2 camera with 4×4 binning. A second camera was used to image mCherry fluorescence for the glycolytic mislocalization experiments. Imaris software (Oxford Instruments) was used to quantify the glucose or amino acid response of cilia and plasma membrane. A custom MATLAB script was used to calculate plasma membrane and ciliary ATP/ADP oscillation parameters including the average amplitude, duty cycle, and frequency of each cell (script available at https://github.com/hrfoster/Merrins-Lab-Matlab-Scripts)^28^.

For cytosolic ATP/ADP and mitochondrial membrane potential measurements, islets from one genotype were alternately barcoded by preincubation with 2 µmol/L DiD (Thermo Fisher Scientific D7757) in 1 mL islet media for 15 min at 37°C to facilitate simultaneous imaging of control and knockout islets. DiD was then imaged using a Cy5 cube (Chroma). For mitochondrial membrane potential measurement, islets were preincubated in 0.83 μM Rhodamine-123 (Sigma) for 5 minutes prior to imaging. Widefield imaging was conducted on a Nikon Ti2 microscope equipped with a 10×/0.5NA SuperFluor objective (Nikon). Excitation was provided by a SOLA SEII 365 (Lumencor) attenuated with an ND4 neutral density filter and set between 5–30% output. Fluorescence emission was collected with a Hamamatsu ORCA-Flash4.0 V2 Digital CMOS camera every 6 s. A single region of interest was used to quantify the average response of each islet using NIS-Elements software (Nikon Instruments). Excitation (x) or emission (m) filters (ET type; Chroma Technology) were used in combination with an FF444/521/608-Di01 dichroic beamsplitter (Semrock) as follows: Rhodamine-123, 500/20x, 535/35m; and Perceval-HR, 430/20x and 500/20x, 535/35m (R500/430).

### Islet perifusion assays

Islet insulin secretion was measured by islet perifusion as previously described^42^. Insulin in the media was assayed using the Promega Lumit Insulin Immunoassay (CS3036A07) and measured using a Tecan Spark Plate Reader.

### Immunohistochemistry

Isolated human and mouse islets were washed with phosphate buffered saline (PBS) and fixed with 4% paraformaldehyde (PFA) for 30 minutes, permeabilized in CAS-Block (Invitrogen) containing 0.3% Triton X-100 for 1 hour at room temperature, and subsequently incubated overnight hours at 4°C with primary antibodies diluted in CAS-Block/Triton. Islets then underwent five 10-minute washes in PBST (PBS with 0.1% Triton X-100) and incubation overnight at 4 °C with secondary antibodies, then washed again. Islets were mounted on glass slides using ProLong Gold Antifade Mountant (Invitrogen) and imaged on a Nikon AXR confocal microscope equipped with NSPARC or a Zeiss Cell Discoverer 7 microscope. Primary antibodies used for cilia immunostaining include: ARL13b (Proteintech 30332-1-AP), AcTUB (Proteintech 66200-1-Ig; Sigma-Aldrich T7451), GLUT1 (Proteintech 21829-1-AP), GLUT2 (Proteintech 20436-1-AP), GCK (Abcam ab88056; Santa Cruz, sc-17819), PFK (Cell Signaling CST 69245), PKm1 (Proteintech 15821-1-AP), PKm2 (Cell Signaling D78A4), LDHA (Cell Signaling C4B5), GAPDH (Cell Signaling D16H11), and GCG (Abcam ab10988). Secondary antibodies were Alexa Fluor goat-anti-mouse and goat-anti-rabbit (Invitrogen) used at 1:500 dilution.

### Immuno-scanning electron microscopy

Intact wildtype mouse islets were adhered to coverslips and fixed overnight in 4% paraformaldehyde in PBS, pH 7.4 at 4°C. Coverslips were then rinsed in PBS three times and transferred into 50 mM glycine in PBS for 15 minutes to quench aldehydes. Following this, samples were first incubated in 1% BSA in PBS for 30 minutes and then labelled with GLUT2 antibody (Proteintech, 20436-1-AP) prepared at dilution 1:200 for 40 hours at 4°C. After washing 5 times for 5 minutes each in 1% BSA in PBS, samples were incubated with anti-rabbit secondary antibody (Jackson Immuno Research, 711-205-152) at 1:20 dilution for 1 hour. Samples were washed 5 times for 5 minutes each in PBS, treated with 2% glutaraldehyde in PBS for 5 minutes and washed again in PBS. Following this, coverslips were incubated with 0.5% osmium tetroxide for 20 minutes on ice and then washed in ultrapure water 3 times for 10 minutes each. Samples were dehydrated in a graded ethanol series and loaded into a critical point drier (Leica EM CPD 300, Vienna, Austria). Coverslips were mounted on aluminum stubs with carbon adhesive tabs and coated with 12 nm of carbon (Leica ACE 600, Vienna, Austria). SEM images were acquired on a FIB-SEM platform (Helios 5 UX DualBeam Fisher Scientific, Brno, Czech Republic) using SEM imaging mode at 5 kV and 0.1 nA.

### Spatial Proteomics of Isolated β-cell Primary Cilia

MIN6 β-cells were cultured in DMEM/high glucose/4 mM glutamine until near-confluence (>90%) and serum-starved for 48 hours to induce ciliation. Deciliation was performed via calcium shock following published protocols^49,60^. Biological triplicates were included. Cells were treated with PBS containing 1 mM EDTA, collected by scraping, and pelleted by centrifugation. The cell pellet was washed with HBSS and resuspended in deciliation solution (20 mM HEPES pH 7.0, 112 mM NaCl, 3.4 mM KCl, 10 mM CaCl_2_, 2.4 mM NaHCO_3_, 20% ethanol) supplemented with 10 μg/mL Cytochalasin D and 1% protease inhibitor cocktail (Sigma-Aldrich). The suspension was incubated for 15 min at 4°C with rotation, then centrifuged at 1,000×g for 5 min at 4°C. The resulting supernatant was serially fractionated by sequential centrifugation: 2,000×g for 30 min at 4°C (cytoplasmic fraction), 10,000×g for 30 min at 4°C (cytoplasmic organelle fraction), followed by 16,000 ×g for 30 min at 4°C (cilia-enriched fraction). Fractions were resuspended in 30 μL PBS containing 30 mM HEPES and protease inhibitor cocktail. Samples were reduced with 4 mM dithiothreitol at 50°C for 30 minutes and subsequently alkylated with 18 mM iodoacetamide for 30 minutes in the dark. Samples were digested with 1 µg of trypsin at 37°C for 16 hours. Following digestion, peptides were acidified with 0.1% trifluoroacetic acid (TFA) and desalted using C18 spin columns. Elution was performed with 50% acetonitrile (ACN) containing 0.1% TFA. The desalted peptides were then lyophilized using a vacuum concentrator and stored at –80°C until used for LC-MS/MS analysis. Peptides were analyzed by LC-MS/MS using a Vanquish Neo UHPLC System coupled to an Orbitrap Eclipse Tribrid Mass Spectrometer with FAIMS Pro Duo interface (Thermo Fisher Scientific). The sample was loaded on a Neo trap cartridge coupled with an analytical column (75 µm ID x 50 cm PepMap^TM^ Neo C18, 2 µm). The flow rate was kept at 250 nL/min. Mobile phase A was 0.1% FA in water, and mobile phase B was 0.1% FA in ACN. The peptides were separated on a 115-minute analytical gradient from 2 to 35% of mobile phase B. For MS acquisition, FAIMS switched between CVs of −35 V and −65 V with cycle times of 1.5 s per CV. MS1 spectra were acquired at 120,000 resolution with a scan range from 375 to 1500 m/z, AGC target set at 300%, and maximum injection time set at Auto mode. Precursors were filtered using monoisotopic peak determination set to peptide, charge state 2 to 7, dynamic exclusion of 60 s with ±10 ppm tolerance. For the MS2 analysis, the isolated ions were fragmented by assisted higher-energy collisional dissociation (HCD) at 30% and acquired in an ion trop. The AGC and maximum IT were Standard and Dynamic modes, respectively. The resulting tandem MS data was queried for protein identification and label-free quantification against the Swiss-Prot *Mus Musculus* database using MaxQuant (version 2.6.7.0). The following search parameters were applied: trypsin digestion with cleavage after K or R (except when followed by P), allowance for up to 2 missed cleavages, carbamidomethylation of cysteine (static modification), and variable modifications including oxidized methionine, deamidated asparagine/glutamine, and protein N-terminal acetylation. Two-tailed t-tests were performed to compare protein abundance between cilia-enriched and cytoplasmic fractions, identifying proteins significantly enriched in the ciliary compartment. Data visualization and pathway enrichment analyses were carried out using WikiPathways and Reactome focusing on glycolysis-associated proteins. The mass spectrometry proteomics data have been deposited to the ProteomeXchange Consortium via the PRIDE^61^ partner repository with the dataset identifier PXD077773 and 10.6019/PXD077773.

## Acknowledgements

The Merrins and Hughes laboratories gratefully acknowledge support from the National Institutes of Health/National Institute of Diabetes and Digestive and Kidney Diseases (R01DK140365) and additional support provided by R01DK113103, R01DK127637, R01DK139640 (to M.J.M.) and R01DK138974 (to J.W.H.). This work was performed with support of the University of Wisconsin Optical Imaging Core, Washington University Center for Cellular Imaging, and the Yale CMC Imaging Core. Mass Spectrometry analyses were performed by the Mass Spectrometry Technology Access Center at the McDonnell Genome Institute (MTAC@MGI) at Washington University School of Medicine, supported by the Diabetes Research Center/NIH grant (P30DK020579), Institute of Clinical and Translational Sciences/NCATS CTSA award (UL1TR002345), and Siteman Cancer Center/NCI CCSG grant (P30CA091842).

## References

1. Landsman, L., Parent, A. & Hebrok, M. Elevated Hedgehog/Gli signaling causes β-cell dedifferentiation in mice. Proceedings of the National Academy of Sciences 108, 17010–17015 (2011).

2. Hughes, J. W. et al. Primary cilia control glucose homeostasis via islet paracrine interactions. Proceedings of the National Academy of Sciences 117, 8912–8923 (2020).

3. Adamson, S. E. & Hughes, J. W. Paracrine signaling by pancreatic islet cilia. Current Opinion in Endocrine and Metabolic Research 35, 100505 (2024).

4. Lau, J. & Hebrok, M. Hedgehog Signaling in Pancreas Epithelium Regulates Embryonic Organ Formation and Adult β-Cell Function. Diabetes 59, 1211–1221 (2010).

5. Volta, F. et al. Glucose homeostasis is regulated by pancreatic β-cell cilia via endosomal EphA-processing. Nat Commun 10, 5686 (2019).

6. Hilgendorf, K. I., Myers, B. R. & Reiter, J. F. Emerging mechanistic understanding of cilia function in cellular signalling. Nat Rev Mol Cell Biol 25, 555–573 (2024).

7. Müller, A. et al. Structure, interaction and nervous connectivity of beta cell primary cilia. Nat Commun 15, 9168 (2024).

8. Sanchez, G. M. et al. The β-cell primary cilium is an autonomous Ca2+ compartment for paracrine GABA signaling. J Cell Biol 222, e202108101 (2022).

9. Wu, C.-T. et al. Discovery of ciliary G protein-coupled receptors regulating pancreatic islet insulin and glucagon secretion. Genes Dev 35, 1243–1255 (2021).

10. Cho, J. H. et al. Islet primary cilia motility controls insulin secretion. Science Advances 8, eabq8486 (2022).

11. Villar, P. S., Delgado, R., Vergara, C., Reyes, J. G. & Bacigalupo, J. Energy Requirements of Odor Transduction in the Chemosensory Cilia of Olfactory Sensory Neurons Rely on Oxidative Phosphorylation and Glycolytic Processing of Extracellular Glucose. J. Neurosci. 37, 5736–5743 (2017).

12. Nunez-Parra, A. et al. Expression and Distribution of Facilitative Glucose (GLUTs) and Monocarboxylate/H+ (MCTs) Transporters in Rat Olfactory Epithelia. Chem Senses 36, 771–780 (2011).

13. Hsu, S. C. & Molday, R. S. Glyceraldehyde-3-phosphate dehydrogenase is a major protein associated with the plasma membrane of retinal photoreceptor outer segments. Journal of Biological Chemistry 265, 13308–13313 (1990).

14. Hsu, S. C. & Molday, R. S. Glycolytic enzymes and a GLUT-1 glucose transporter in the outer segments of rod and cone photoreceptor cells. Journal of Biological Chemistry 266, 21745–21752 (1991).

15. Lopez-Escalera, R., Li, X. B., Szerencsei, R. T. & Schnetkamp, P. P. M. Glycolysis and glucose uptake in intact outer segments isolated from bovine retinal rods. Biochemistry 30, 8970–8976 (1991).

16. Narita, K., Nagatomo, H., Kozuka-Hata, H., Oyama, M. & Takeda, S. Discovery of a Vertebrate-Specific Factor that Processes Flagellar Glycolytic Enolase during Motile Ciliogenesis. iScience 23, (2020).

17. Pazour, G. J., Agrin, N., Leszyk, J. & Witman, G. B. Proteomic analysis of a eukaryotic cilium. J Cell Biol 170, 103–113 (2005).

18. Mitchell, B. F., Pedersen, L. B., Feely, M., Rosenbaum, J. L. & Mitchell, D. R. ATP production in Chlamydomonas reinhardtii flagella by glycolytic enzymes. Mol Biol Cell 16, 4509–4518 (2005).

19. Merrins, M. J., Corkey, B. E., Kibbey, R. G. & Prentki, M. Metabolic cycles and signals for insulin secretion. Cell Metabolism 34, 947–968 (2022).

20. Deeney, J. T., Prentki, M. & Corkey, B. E. Metabolic control of β-cell function. Seminars in Cell & Developmental Biology 11, 267–275 (2000).

21. Matschinsky, F. M. & Ellerman, J. E. Metabolism of Glucose in the Islets of Langerhans. Journal of Biological Chemistry 243, 2730–2736 (1968).

22. Fridlyand, L. E. & Philipson, L. H. Mechanisms of glucose sensing in the pancreatic β-cell. Islets 3, 224–230 (2011).

23. Maechler, P., Wang, H. & Wollheim, C. B. Continuous monitoring of ATP levels in living insulin secreting cells expressing cytosolic firefly luciferase. FEBS Letters 422, 328–332 (1998).

24. Ainscow, E. K. & Rutter, G. A. Glucose-Stimulated Oscillations in Free Cytosolic ATP Concentration Imaged in Single Islet β-Cells: Evidence for a Ca2+-Dependent Mechanism. Diabetes 51, S162–S170 (2002).

25. Ainscow, E. K. & Brand, M. D. Top-down control analysis of ATP turnover, glycolysis and oxidative phosphorylation in rat hepatocytes. European Journal of Biochemistry 263, 671–685 (1999).

26. Fridlyand, L. E., Ma, L. & Philipson, L. H. Adenine nucleotide regulation in pancreatic β-cells: modeling of ATP/ADP-Ca2+ interactions. American Journal of Physiology-Endocrinology and Metabolism 289, E839–E848 (2005).

27. Lewandowski, S. L. et al. Pyruvate Kinase Controls Signal Strength in the Insulin Secretory Pathway. Cell Metabolism 32, 736–750.e5 (2020).

28. Foster, H. R. et al. β-cell deletion of the PKm1 and PKm2 isoforms of pyruvate kinase in mice reveals their essential role as nutrient sensors for the KATP channel. eLife 11, e79422 (2022).

29. Ho, T., Potapenko, E., Davis, D. B. & Merrins, M. J. A plasma membrane-associated glycolytic metabolon is functionally coupled to KATP channels in pancreatic α and β cells from humans and mice. Cell Reports 42, (2023).

30. Wang, H. et al. Organization of a functional glycolytic metabolon on mitochondria for metabolic efficiency. Nat Metab 6, 1712–1735 (2024).

31. Jang, S. et al. Glycolytic Enzymes Localize to Synapses under Energy Stress to Support Synaptic Function. Neuron 90, 278–291 (2016).

32. Sepp, M. et al. Tight coupling of Na+/K+-ATPase with glycolysis demonstrated in permeabilized rat cardiomyocytes. PLoS One 9, e99413 (2014).

33. Xu, K. Y., Zweier, J. L. & Becker, L. C. Functional coupling between glycolysis and sarcoplasmic reticulum Ca2+ transport. Circ Res 77, 88–97 (1995).

34. Korge, P. & Campbell, K. B. Local ATP regeneration is important for sarcoplasmic reticulum Ca2+ pump function. Am J Physiol 267, C357–366 (1994).

35. Weiss, J. N. & Lamp, S. T. Glycolysis preferentially inhibits ATP-sensitive K+ channels in isolated guinea pig cardiac myocytes. Science 238, 67–69 (1987).

36. Tantama, M., Martínez-François, J. R., Mongeon, R. & Yellen, G. Imaging energy status in live cells with a fluorescent biosensor of the intracellular ATP-to-ADP ratio. Nature Communications 4, 2550 (2013).

37. Su, S. et al. Genetically encoded calcium indicator illuminates calcium dynamics in primary cilia. Nature Methods 10, 1105–1107 (2013).

38. Benninger, R. K. P., Zhang, M., Head, W. S., Satin, L. S. & Piston, D. W. Gap Junction Coupling and Calcium Waves in the Pancreatic Islet. Biophys J 95, 5048–5061 (2008).

39. Polino, A. J., Sviben, S., Melena, I., Piston, D. W. & Hughes, J. W. Scanning electron microscopy of human islet cilia. Proceedings of the National Academy of Sciences 120, e2302624120 (2023).

40. DiGruccio, M. R. et al. Comprehensive alpha, beta and delta cell transcriptomes reveal that ghrelin selectively activates delta cells and promotes somatostatin release from pancreatic islets. Mol Metab 5, 449–458 (2016).

41. Sekine, N. et al. Low lactate dehydrogenase and high mitochondrial glycerol phosphate dehydrogenase in pancreatic beta-cells. Potential role in nutrient sensing. J. Biol. Chem. 269, 4895–4902 (1994).

42. Jin, E. et al. Amino Acid Sensing by the α-Cell Mitochondrial Phosphoenolpyruvate Cycle Regulates Intracellular Ca2+ Levels Without Affecting Glucagon Secretion. Diabetes 75, 483–493 (2026).

43. Li, C. et al. Regulation of Leucine-stimulated Insulin Secretion and Glutamine Metabolism in Isolated Rat Islets*. Journal of Biological Chemistry 278, 2853–2858 (2003).

44. Sener, A. & Malaisse, W. J. L-leucine and a nonmetabolized analogue activate pancreatic islet glutamate dehydrogenase. Nature 288, 187–189 (1980).

45. Gylfe, E. Comparison of the effects of leucines, non-metabolizable leucine analogues and other insulin secretagogues on the activity of glutamate dehydrogenase. Acta Diabetol Lat 13, 20–24 (1976).

46. Hu, Q. & Nelson, W. J. Ciliary diffusion barrier: The gatekeeper for the primary cilium compartment. Cytoskeleton 68, 313–324 (2011).

47. Hu, Q. et al. A Septin Diffusion Barrier at the Base of the Primary Cilium Maintains Ciliary Membrane Protein Distribution. Science 329, 436–439 (2010).

48. Merrins, M. J. & Kibbey, R. G. Glucose Regulation of β-Cell KATP Channels: It Is Time for a New Model! Diabetes 73, 856–863 (2024).

49. Steidl, M. E. et al. Primary cilia sense glutamine availability and respond via asparagine synthetase. Nat Metab 5, 385–397 (2023).

50. van der Meulen, T. et al. Virgin Beta Cells Persist throughout Life at a Neogenic Niche within Pancreatic Islets. Cell Metabolism 25, 911–926.e6 (2017).

51. Thorens, B. et al. Ins1(Cre) knock-in mice for beta cell-specific gene recombination. Diabetologia 58, 558–565 (2015).

52. Davidson, S. M. et al. Pyruvate Kinase M1 Suppresses Development and Progression of Prostate Adenocarcinoma. Cancer Res 82, 2403–2416 (2022).

53. Li, Q. et al. PKM1 Exerts Critical Roles in Cardiac Remodeling Under Pressure Overload in the Heart. Circulation 144, 712–727 (2021).

54. Israelsen, W. J. et al. PKM2 Isoform-Specific Deletion Reveals a Differential Requirement for Pyruvate Kinase in Tumor Cells. Cell 155, 397–409 (2013).

55. Gregg, T. et al. Pancreatic β-Cells From Mice Offset Age-Associated Mitochondrial Deficiency With Reduced KATP Channel Activity. Diabetes 65, 2700–2710 (2016).

56. Merrins, M. J. et al. Phase Analysis of Metabolic Oscillations and Membrane Potential in Pancreatic Islet β-Cells. Biophysical Journal 110, 691–699 (2016).

57. Koberstein, J. N. et al. Monitoring glycolytic dynamics in single cells using a fluorescent biosensor for fructose 1,6-bisphosphate. Proceedings of the National Academy of Sciences 119, e2204407119 (2022).

58. Gold, M. G. et al. Architecture and dynamics of an A-kinase anchoring protein 79 (AKAP79) signaling complex. Proceedings of the National Academy of Sciences 108, 6426–6431 (2011).

59. Tenner, B. et al. Spatially compartmentalized phase regulation of a Ca2+-cAMP-PKA oscillatory circuit. eLife 9, e55013 (2020).

60. Ishikawa, H., Thompson, J., Yates, J. R. & Marshall, W. F. Proteomic Analysis of Mammalian Primary Cilia. Current Biology 22, 414–419 (2012).

61. Perez-Riverol, Y. et al. The PRIDE database at 20 years: 2025 update. Nucleic Acids Res 53, D543–D553 (2025).

