## Supplemental Figures for "Compartmentalized glycolysis powers ATP production in primary cilia and engages mitochondria via the phosphoenolpyruvate cycle"

**Fig. S1: Primary cilia are autonomous metabolic compartments with distinct ATP/ADP dynamics from the plasma membrane in mouse islet β-cells.**
**A–C**, Quantification of ATP/ADP oscillatory parameters measured using $5HT_6$-Perceval-HR, including amplitude (**A**), frequency (**B**), and duty cycle (**C**), plotted at the level of individual β-cell. Each point represents a single cilium or plasma membrane, with bars indicating mean ± SEM. Data correspond to the same experiment shown in Fig. 1B–G but are presented without islet-level averaging to illustrate cell-to-cell variability within islets (*n* = 260 β-cells from 26 islets across 3 mice).

**Fig. S2: Validation of glucose transporter and glycolytic enzyme localization to β-cell primary cilia.**
**A,** Three-dimensional immuno-scanning electron microscopy (immuno-SEM) showing surface GLUT2 immunogold labeling on mouse β-cells. GLUT2 signal is detected on primary cilia. No signal was detected in secondary antibody–only controls. Scale bar, 200 nm. **B,** Immunofluorescence images of mouse islets showing GLUT2 (red), acetylated α-tubulin (AcTUB, green), and nuclei (DAPI, blue). Insets highlight GLUT2 localization within primary cilia marked by acetylated α-tubulin. Scale bar, 1 μm. **C,D,** Immunofluorescence validation of pyruvate kinase isoform localization in mouse islet cilia. PKm1 (**C**) and PKm2 (**D**) (magenta) are shown with the ciliary marker Arl13b (green) and glucagon (GCG, red). Arrows indicate primary cilia. Images from control (Con) and β-cell–specific knockout (βKO) islets confirm antibody specificity. Scale bar, 5 μm. **E,** Localization of lactate dehydrogenase A (LDHA, magenta) in primary cilia of mouse and human islets, co-stained with ARL13B (green). Arrows indicate primary cilia. Scale bar, 5 μm. **F,** Immunoblot analysis of acetylated α-tubulin and cyclophilin A in isolated ciliary and non-ciliary fractions. Acetylated α-tubulin is enriched in the ciliary fraction, whereas cyclophilin A is predominantly detected in the non-ciliary (cytosolic) fraction, confirming cilia purification. Quantification of acetylated α-tubulin abundance is shown at right.

**Fig. S3: Workload-dependent ATP/ADP dynamics in β-cell cilia and plasma membrane.**
**A,** Proposed model of compartmentalized glycolysis and ATP production in the primary cilium, highlighting the role of pyruvate kinase (PK). **B,** ATP/ADP dynamics at the plasma membrane and cilia during glucose stimulation from 2 mM glucose (2G) to 10 mM glucose (10G) in the presence or absence of cyclopiazonic acid (CPA, 50 μM), an inhibitor of the sarco/endoplasmic reticulum $Ca^{2+}$-ATPase (SERCA), followed by diazoxide (Dz, 200 μM) treatment. Traces show average responses, with quantification of the area under the curve (AUC) at right (vehicle, *n* = 324 cilia from 27 islets across 3 mice; CPA, *n* = 420 cilia from 41 islets across 3 mice). **C,D,** Glucose-stimulated insulin secretion in control and PKm1-βKO (**C**) or PKm2-βKO (**D**) islets. Data are quantified as AUC from PKm1-βKO (*n* = 10 mice) and controls (*n* = 10 mice); PKm2-βKO (*n* = 5 mice) and controls (*n* = 5 mice).

**Fig. S4: Additional characterization of metabolic and functional phenotypes of PCK2-βKO and PKm1-βKO.**

**A,** Glucose-stimulated insulin secretion in control (Con) and PCK2-βKO islets. Data are quantified as AUC from PCK2-βKO (*n* = 15 mice) and controls (*n* = 15 mice). **B,** Mitochondrial membrane potential (ΔΨm) measured using Rhodamine-123 in control (Con) and PKm1-βKO islets during stimulation with 10 mM glucose (10G), followed by cyanide (KCN) to induce mitochondrial depolarization. ΔΨm was normalized to fluorescence after KCN-induced depolarization, and quantified as AUC for 10G treatment from PKm1-βKO (*n* = 97 islets from 3 mice) and littermate controls (*n* = 99 islets from 3 mice).

**Fig. S5: Supporting functional and imaging analyses of PCK2-βKO and PKm1-βKO.**

**A,B,** Insulin secretion in control (Con) and PCK2-βKO (**A**) or PKm1-βKO (**B**) islets in response to leucine (1.5 mM) stimulation in the presence of 0.5 mM glutamine. Data are quantified as AUC from PCK2-βKO (*n* = 6 mice) and controls (*n* = 6 mice); PKm1-βKO (*n* = 3 mice) and controls (*n* = 3 mice). **C,** Representative three-dimensional images of mouse islets expressing cilia-targeted biosensors for ATP/ADP ($5HT_6$-PercevalHR), pyruvate ($5HT_6$-PyronicSF), and fructose-1,6-bisphosphate (FBP; $5HT_6$-HYLight). Insets show higher magnification views of the boxed regions, highlighting ciliary and plasma membrane localization of each biosensor. Scale bars, 50 μm.

### Supplemental Figure 1

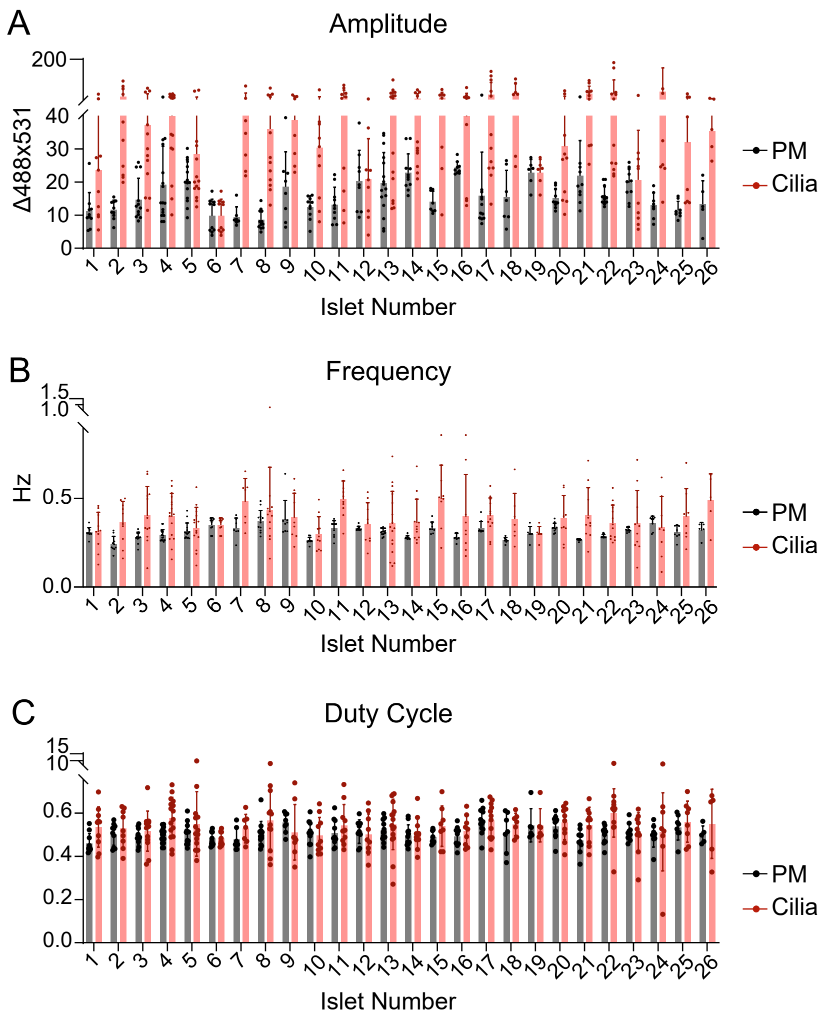

### Supplemental Figure 2

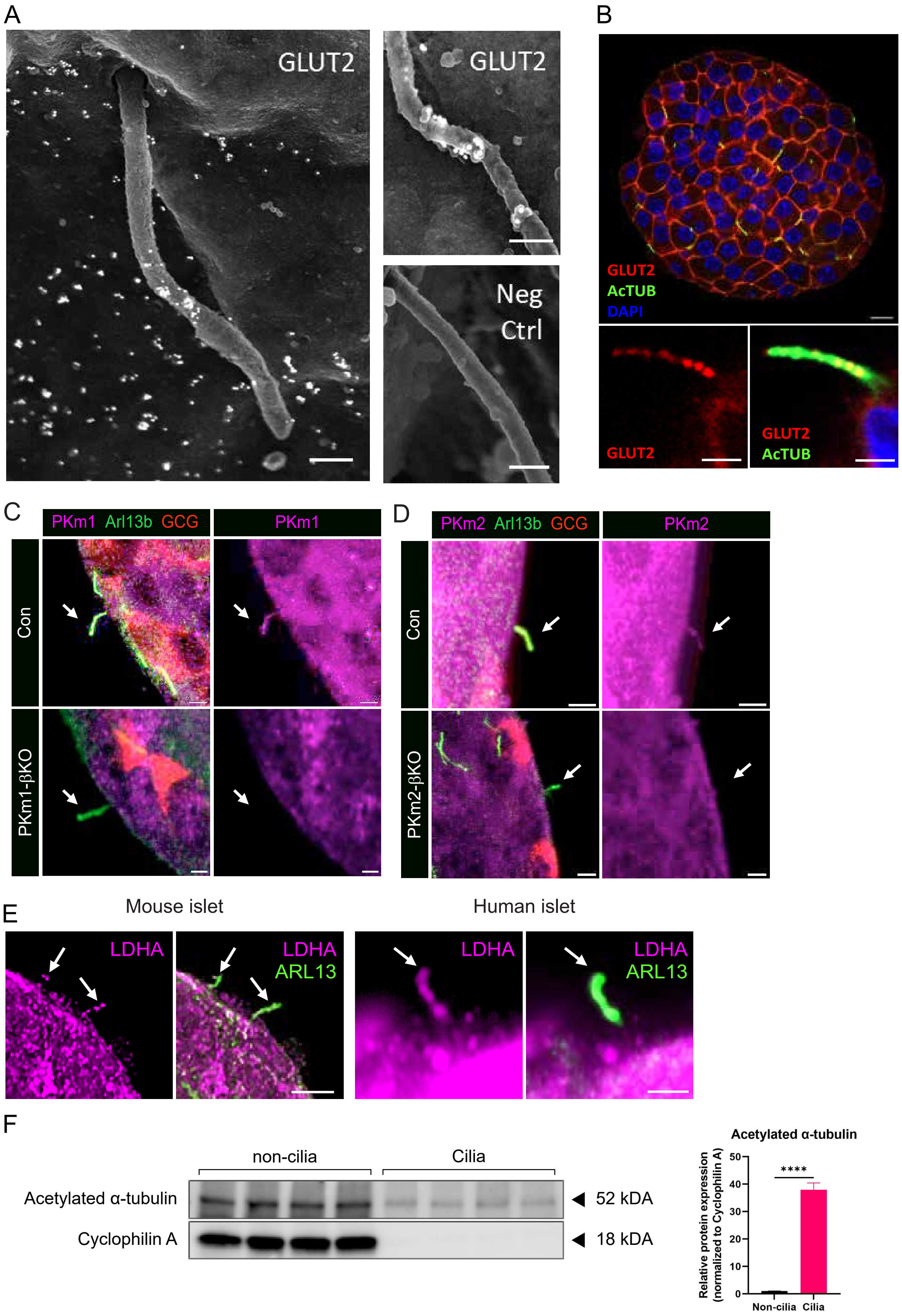

Supplemental Figure 3

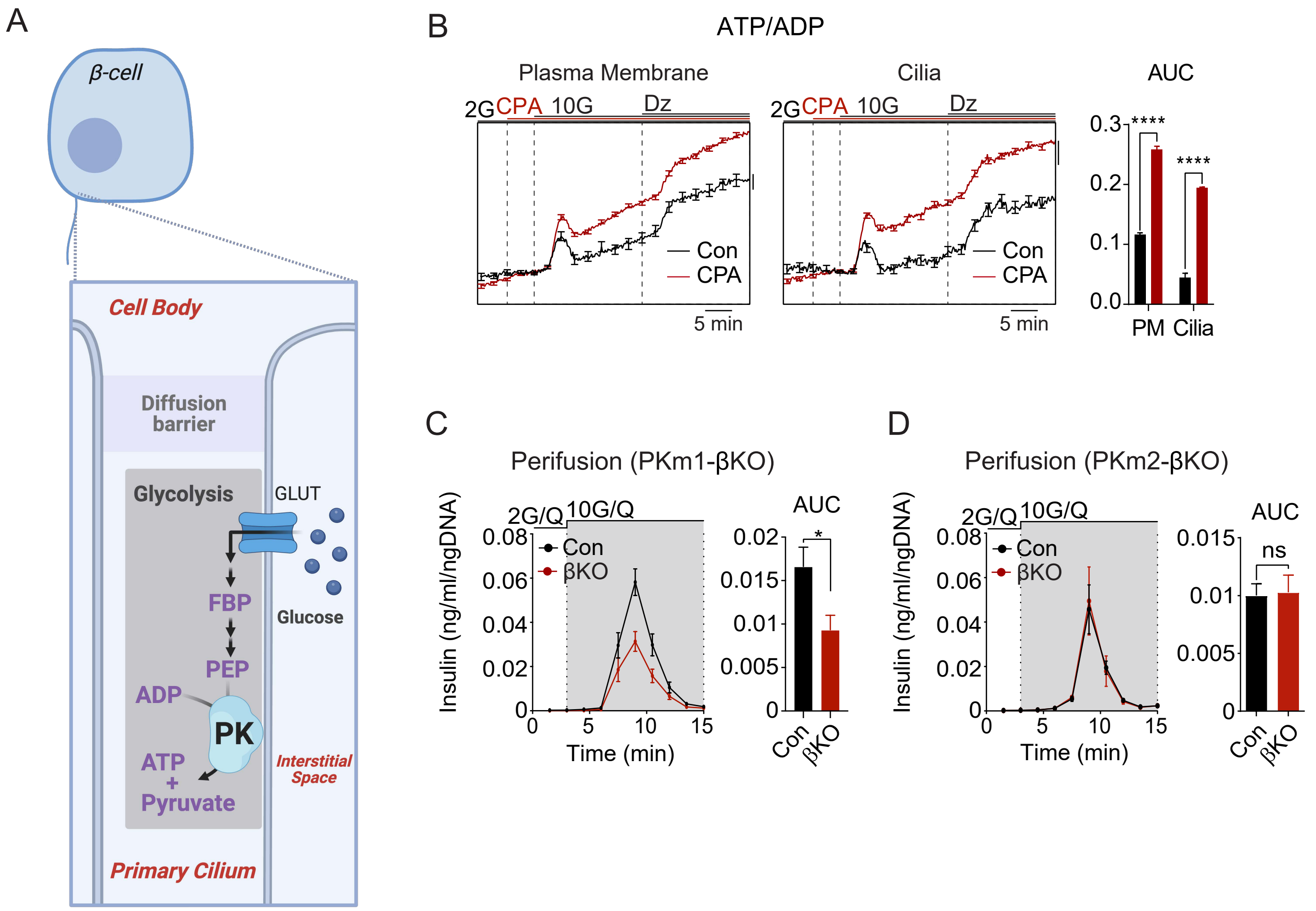

### Supplemental Figure 4

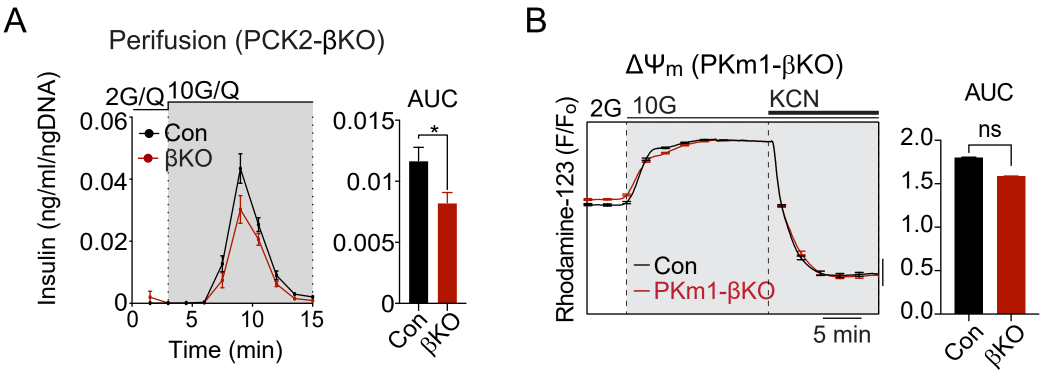

#### Supplemental Figure 5

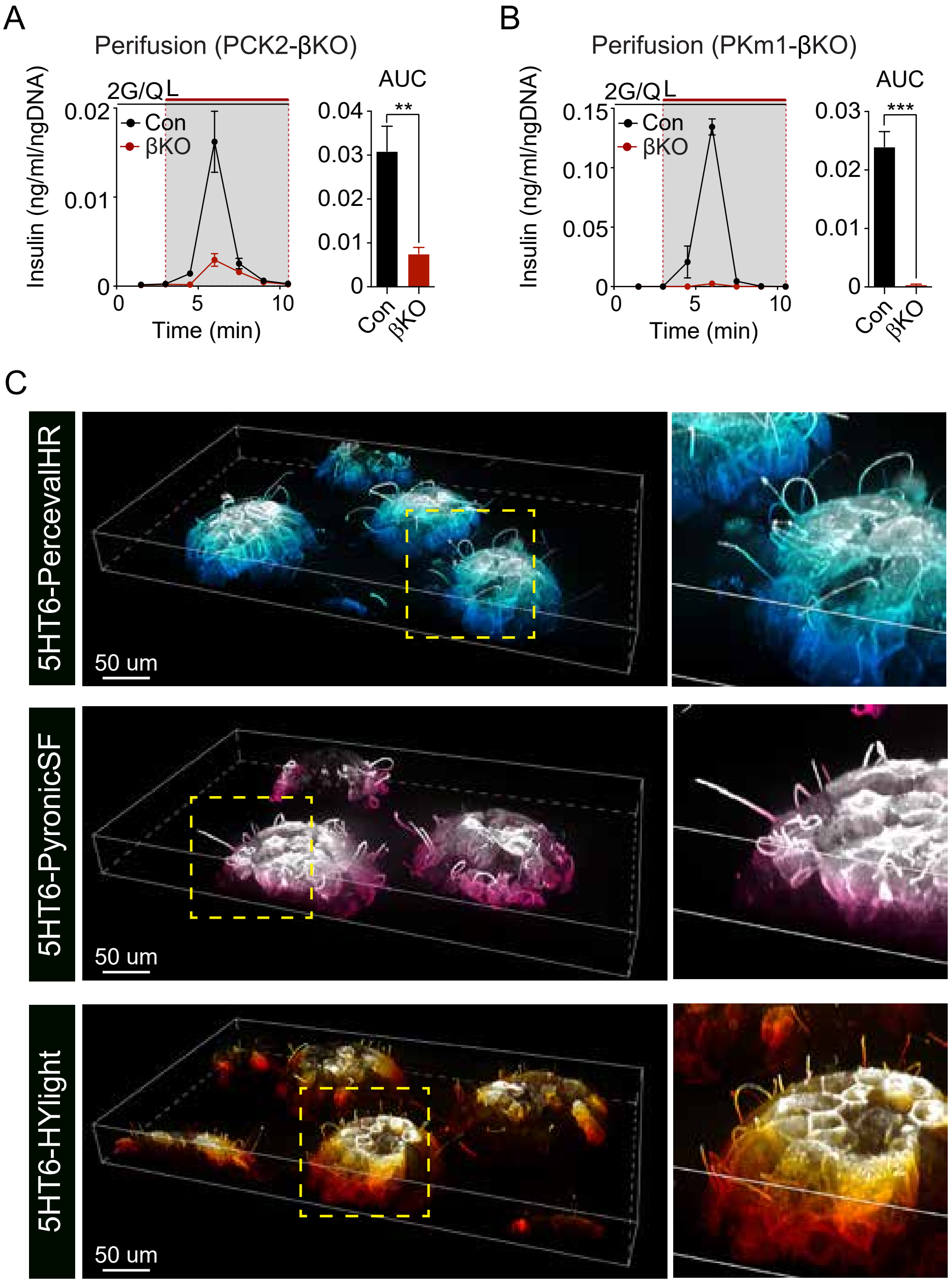
